# Monitoring the poroelastic response of collagen gels with embedded senescent fibroblasts reveals tissue softening associated with collagen network reorganization

**DOI:** 10.1101/2025.04.18.649513

**Authors:** Jean Cacheux, Thomas Quénan, Daniel Alcaide, Jose Ordonez-Miranda, Laurent Jalabert, Shizuka Nakano, Makoto Nakanishi, Pierre Cordelier, Aurélien Bancaud, Yukiko T. Matsunaga

## Abstract

The poroelastic properties of tissues regulate molecular transport and mechanical signaling, yet their evolution during aging remains poorly understood. In particular, senescent fibroblasts accumulate in aged tissues, contributing to extracellular matrix (ECM) remodeling, but their impact on tissue mechanics and permeability is unclear. In this study, we developed a microfluidic-based in vitro model to assess the poroelastic properties of collagen gels embedded with senescent fibroblasts over time. Our approach integrates periodic pressure actuation with real-time pressure monitoring in a sealed air cavity, enabling the detection of fluid permeation and solid matrix deformations. We analyze our data using analytical and numerical models based on a porohyperelastic framework. This framework combines compressible Neo-Hookean elasticity with the Kozeny–Carman permeability relationship. We demonstrate that senescent fibroblasts induce a progressive softening of the ECM without altering its permeability. Immunostaining reveals that this softening correlates with structural reorganization of the collagen network, characterized by increased branching and network remodeling. Our findings provide insights into the biomechanical effects of senescent fibroblasts on ECM homeostasis. We further argue that our platform offers a unique solution to investigate ECM remodeling not only in aging but also fibrosis, cancer progression, or regenerative medicine strategies.

## Introduction

Tissues serve as porous solid matrices through which interstitial fluid flows, transporting oxygen and nutrients from the bloodstream to the cells, while concurrently allowing the removal of cellular waste products. The intricate fluid dynamics between blood and tissues have been described by the Starling principle for more than a century. The textbook model delineates a balance between hydrostatic and oncotic pressures, with the latter arising from the protein component of osmotic pressure [1,2]. It has recently been updated to include the lymphatic system, which plays a crucial role in fluid drainage within tissues [3]. The connective tissue, a complex material composed of the ECM (fiber proteins, proteoglycans, *etc*.) and stromal cells such as fibroblasts, contributes to molecular exchanges, in particular through its microstructural properties that set the rate of fluid flows.

The physical properties of connective tissues are highly dynamic and evolve throughout their lifetime [4]. For example, aging in human tissues leads to microstructural changes that typically result in mechanical softening, as observed in the skin [5]. This softening is primarily attributed to a decrease in the biosynthesis of precollagen 1 and the subsequent reduction in collagen fibril concentration [6], which are the most abundant components of the ECM [7]. Additionally, the expression of various metalloproteinases in aged human dermal skin [8] accelerates collagen fibril fragmentation and elastin degradation [9,10], further contributing to ECM breakdown. The combination of diminished collagen production and increased degradation underlies the progressive softening of aged skin. However, evidence also indicates excessive cross-linking of collagen fibers during skin aging [11], notably due to sun exposure [12]. This cross-linking, which is expected to stiffen the ECM, presents a seemingly contradictory mechanism in the context of skin softening. These findings suggest that multiple, potentially opposing processes — softening through degradation and stiffening via cross-linking — jointly influence ECM properties during aging.

Senescent cells emerge as key players, accumulating over time and significantly contributing to age-related diseases. Interestingly, their suppression has been demonstrated to extend lifespans in animal models [13–15]. When present in the tissue, they induce a state of chronic inflammation, leading to multiple biological events, such as high levels of metalloproteinase, high concentrations of reactive oxygen species, or elevated levels of elastase activity that might compromise the integrity of the ECM networks [16]. Senescent cells have been thoroughly investigated *in vitro*, showing these cells could either exert excessive traction stress [17] or lead to a loss of traction force [18]. Yet, to the best of our knowledge, the elasticity of tissues embedded with senescent fibroblasts has not been assayed nor have the implications of aging on tissue permeation properties been explored. It is therefore unclear whether and to what extent senescent cells contribute to the homeostasis of molecular exchanges in tissues.

We postulate that these questions remain debated because assessing the poroelastic properties of soft tissues in living organisms over extended periods is challenging. Other techniques provide only a partial mechanical analysis and risk altering the sample due to stress induced by solid probes, as seen in gold-standard indentation methods. In this report, we use a tissue-engineering approach to fabricate an elementary *in vitro* tissue model-on-a-chip, specifically designed to investigate the impact of senescent dermal fibroblasts on collagen gels. Note that senescent fibroblasts do not proliferate, allowing us to provide readouts of their interaction with the ECM unbiased by cell growth dynamics. We present a contact-free technology based on periodic fluid injection and pressure measurements in an air cavity downstream of this injection to dynamically monitor tissue elasticity and permeability over time. We support experimental data with extensive analytical and finite-element models, which include non-linear mechanical and hydrodynamical (poro-hyperelastic) properties. We specifically integrate the compressible Neo-Hookean stress-strain mechanical model [19] with the normalized Kozeny–Carman permeability model [20], and fit and monitor the response of the collagen extracellular matrix (ECM). We establish that senescent fibroblasts soften the ECM without changing its permeability, and support this observation with immunostainings that prove the structural rearrangements of the collagen network induced by senescent fibroblasts. We finally discuss the implications of these observations in the context of skin aging.

## Results

### Experimental set up

We investigated the effects of senescent skin fibroblasts on collagen gels using a silicone microfluidic system [17]. This microfluidic chip consisted of central cavity of 30 µL, in which unreticulated collagen gels are poured (left panel in Fig. 1A). It also served as scaffold to introduce an acupuncture needle of 200 µm in diameter (right panel in Fig. 1A). Last, the chip contained two smaller lateral cavities of 7 µL that were used as reservoirs to inject fluid in the lumen left by the acupuncture needle. The fabrication of the ECM was performed by mixing senescent fibroblasts at a concentration of 200 cells/µL in an unreticulated collagen solution at 2.4 mg/mL (see details in the methods section). The cell concentration was chosen to approximate the physiological range observed in human dermis, where senescent fibroblasts constitute up to 1–15% of total fibroblasts—estimated at approximately 5,500 cells/µL in native dermis [21]. This solution was poured in the central reservoir of the microfluidic, the acupuncture needle being already inserted into its sheath. After reticulation at 37°C, the needle was removed in order to imprint the lumen of 200 µm. Note that ECM was covalently attached to the bottom and lateral surfaces of the microfluidic chip. We finally poured 1 mL of medium on top of the ECM, and cultured it during 10 days with medium change every other day.

**Figure 1:**
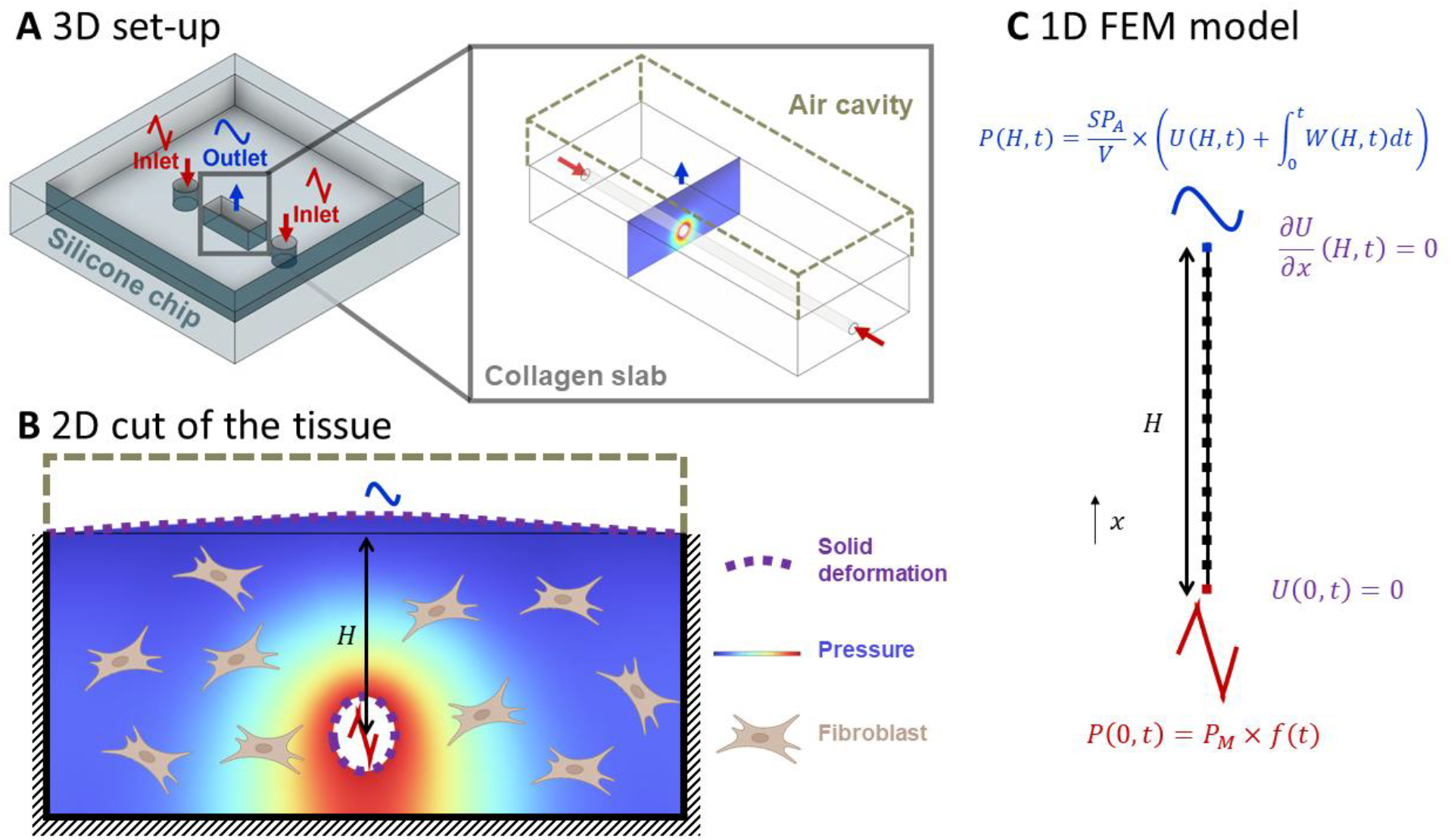
Design of the study and principle of the characterization technique. **A)** The setup consists of a silicone chip with two inlets pointed in red that allow the periodic injection of fluid in the ECM. These inlets are connected to the central cavity with the ECM imprinted with a hollow tube. The top surface of the tissue interfaces with an air cavity (dashed brown lines), where the outlet pressure (in blue) is monitored. **B)** When pressure is applied to the tissue, it diffuses in the material and induces both liquid flow and solid deformation. These mechanisms collectively compress the air in the sealed cavity. **C)** These mechanisms are approximated using a 1D model. The inlet is assumed to be mechanically clamped and subjected to triangular wave periodic pressure, while the pressure at the outlet, *x = H*, is proportional to the integral of the permeation flow *W* through the outlet surface over time and the solid deformation *U*. The outlet is treated as a free surface for the solid deformation.

The microfluidic chip was also used as support to perform poromechanical characterizations, using a variation of the methodology published elsewhere [22]. Briefly, we used 3D printing to fabricate a device that allowed us to separate the lateral injection channels from the ECM, that were considered as the inlets and outlet, respectively (left panel of Fig. 1A). The outlet was sealed, leaving an air cavity of volume *V* = 2 mL (Fig. 1A-B), in which the pressure was monitored in real time. The entire system was placed in a thermally insulated enclosure to measure changes in the volume of the air cavity through pressure measurements, assuming ideal gas behavior. Indeed, as we recently showed [22], fluid injection in the tissue induces poroelastic deformation of the solid and permeation flow, both leading to the compression of the air cavity (Fig. 1B, see details in the next section). These two effects occur on separate time scales, allowing the determination of the elasticity and the permeability using one single kinetic measurement [22]. In this report, we employed periodic triangular wave pressure actuation, as this signal offers a broad spectrum of harmonic frequencies, enabling more comprehensive sampling of the ECM properties. This approach aligns with the recent work from Fiori *et al*., who demonstrated that periodic loading in soft porous materials induces coupled deformation and flow, offering insights into the interplay of mechanical and fluidic responses [23].

### Description of the model and analytical solution in the linear poroelastic regime

The geometry of our system shares some similarities with that of a porous elastic cylinder, in which fluid is injected, as described by Kenyon. [24] However, the fluid can only escape in the vertical direction, toward the air cavity, due to the boundary conditions (Fig. 1B). We previously demonstrated that this vertically confined response can be described using a 1D poroelastic model, showing excellent agreement with 2D numerical simulations [22]. We therefore adopt this strategy to derive an analytical solution for the pressure in the air cavity. Taking the formalism of Kenyon for linear poroelastic materials, two parameters are necessary to describe the poroelastic response. First, we define the permeability *κ* with Darcy’s law

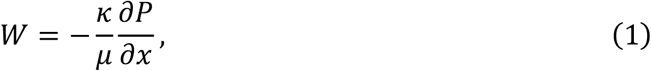

where *µ* is the fluid viscosity, *W* the permeation flow velocity, and ∂*P*⁄∂*x* the pressure field gradient. The pressure gradient also induces the deformation of the solid *U*(*x, t*), and the mechanical equilibrium yields:

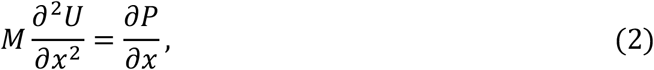

with *M* being the P-wave elastic modulus of the material. Because the total rate of liquid and solid transport is conservative (*W* + ∂*U*⁄∂*t* = *A*(*t*), where *A*(*t*) is a constant of integration), the combination of Eqs. 1 and 2 determines the spatial distribution and temporal evolution of the pressure and deformation fields. These equations are solved with the boundary conditions indicated in Fig. 1C of no displacement at the bottom (*x* = 0) due to the attachment of the ECM to the walls, and no contact stress at the outlet (*x* = *H*). The inlet pressure is a triangular signal represented as a sum of harmonic sine functions, with *ω*_0_ being the fundamental pulsation. The outlet pressure (*i*.*e*., in the air cavity) is dictated by the integral of the filtration velocity *W* over time and the solid deformation *U*.

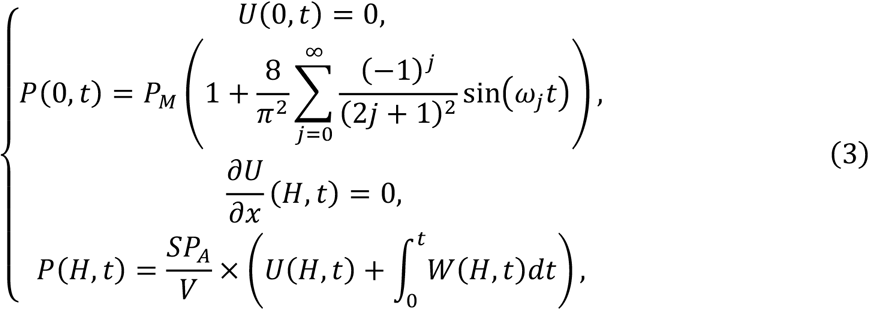

where *ω*_*j*_ = (2*j* + 1)*ω*_0_ with *ω*_0_ = 2*π*⁄*T*_0_, the cross section *S* = 15 mm^2^ and the height of the tissue *H* = 0.8 mm. Considering that the relative volume changes in the cavity are small, on the order of 10^−3^, we can use the ideal gas approximation. We can hence solve this set of Eqs. 3 using the Fourier transform (see Supplementary Materials for details of the calculation), and extract the pressure in the cavity,

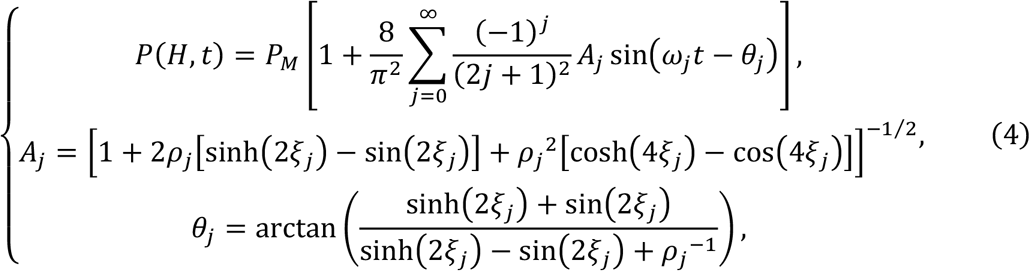

where 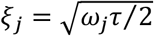 and *ρ*_*j*_ = *αξ*_*j*_⁄(cosh(2*ξ*_*j*_) + cos (2*ξ*_*j*_)) involve the intrinsic constants *α* = *VM*⁄*SHP*_*A*_ and *τ* = *µH*^2^⁄*kM*. The same constants appear when we solve for the response to a step function in pressure [22]. We explained their physical meaning by considering the model situation of an open-air cavity *P*(*H, t*) = 0. At the steady state, we have *U*(*H*) = *P*_*M*_*H*⁄2*M* and *W* = *k P*_*M*_⁄*µH*. The characteristic time *τ* = 2*U*(*H*)⁄*W* thus represents the ratio between twice the displacement and the filtration velocity, while *α* is the ratio of the inlet pressure to the pressure change induced by the solid deformation. Further, we note that this analytical model exactly matched the results of linear finite element simulations (see methods and Supplementary Fig. S1).

### Finite element model in the non-linear poroelastic regime

Collagen gels are known to exhibit non-linear properties in both elasticity [25] and permeability [22]. Since we used triangular periodic pressure actuation, which triggers responses at multiple timescales, we decided to develop a non-linear poroelastic model. The model was parametrized based on the local volume change of the solid matrix, characterized using the Jacobian determinant *J* = 1/(1 − ∂*U*⁄∂*x*) [26]. To represent the relationship between permeability and solid deformation, we employed the widely accepted Kozeny–Carman model, following approaches such as those of MacMinn *et al*. [27],

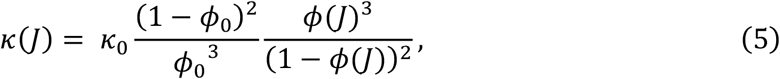

where *ϕ*_0_ and *κ*_0_ are the porosity and the permeability at rest, and *ϕ*(*J*) is the local porosity due to the deformation. Note that we assume that the solid and fluid phases are individually incompressible, implying that *ϕ*(*J*) can be define as follows [26],

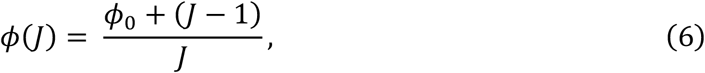

For highly hydrated collagen gels, such as the low concentration collagen gels that are used in this study, *ϕ*_0_ tends to 1, the Taylor expansion of Eq. 5 is

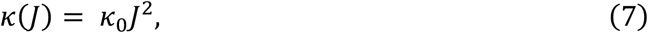

Notably, this quadratic phenomenological relationship between the permeability and the solid deformation is in agreement with the response described by Castro *et al*. [28], where the authors fitted a power-law parameter in the range of 1.8 to 3.5 for collagen gel concentrations from 2 to 4 mg/mL.

Next, the mechanical properties were modelled with the compressible neo-Hookean model, which is widely used in engineering to describe the response of hyperelastic materials, and particularly soft biological tissues [29]. We parametrized this model with the incremental Young modulus and Poisson ratio, as calculated by Scott [30],

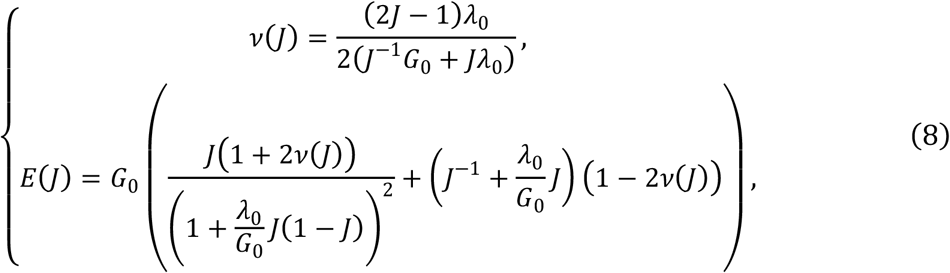

Where *G*_0_ and *λ*_0_ are the shear modulus and Lamé’s first parameter of the material, respectively. In this non-linear regime, we used finite element simulations with the COMSOL Multiphysics software (see Methods). For the sake of simplicity, and to ensure a large screening of the parameter space (*G*_0_, *λ*_0_, *κ*_0_), the model has been developed in 1D, considering a homogeneous poroelastic material, in which the fluid and solid displacements are unidirectional.

### Calibration and optimization of the platform with cross-linked collagen gels

We started by examining the behavior of chemically cross-linked collagen gels under a periodic triangular inlet pressure. This choice was motivated by the material’s key properties: a constant permeability irrespective of the applied pressure and a symmetrical elastic response in both tension and compression [22]. These characteristics indeed justify the use of the linear poroelastic model. For an imposed inlet pressure *P*_*M*_ of 500 Pa, the collagen gel was submitted to a pressure variation from - 500 Pa to 500 Pa, corresponding to a pressure magnitude of 1000 Pa at quasi-steady state, and the period *T*_0_ of the triangle was modulated from 7.4 to 23.6 s (green dashed line in Fig. 2A). The outlet pressure was precisely centered at *P*_*M*_ across all three periods, with its amplitude more than doubling as the period increased from 7.4 to 23.6 s (circles in Fig. 2A). This increase can be readily attributed to the contribution of the permeation flow (see Eq. (3)), an integral term that grows as the pressure gradient is sustained for longer durations.

**Figure 2:**
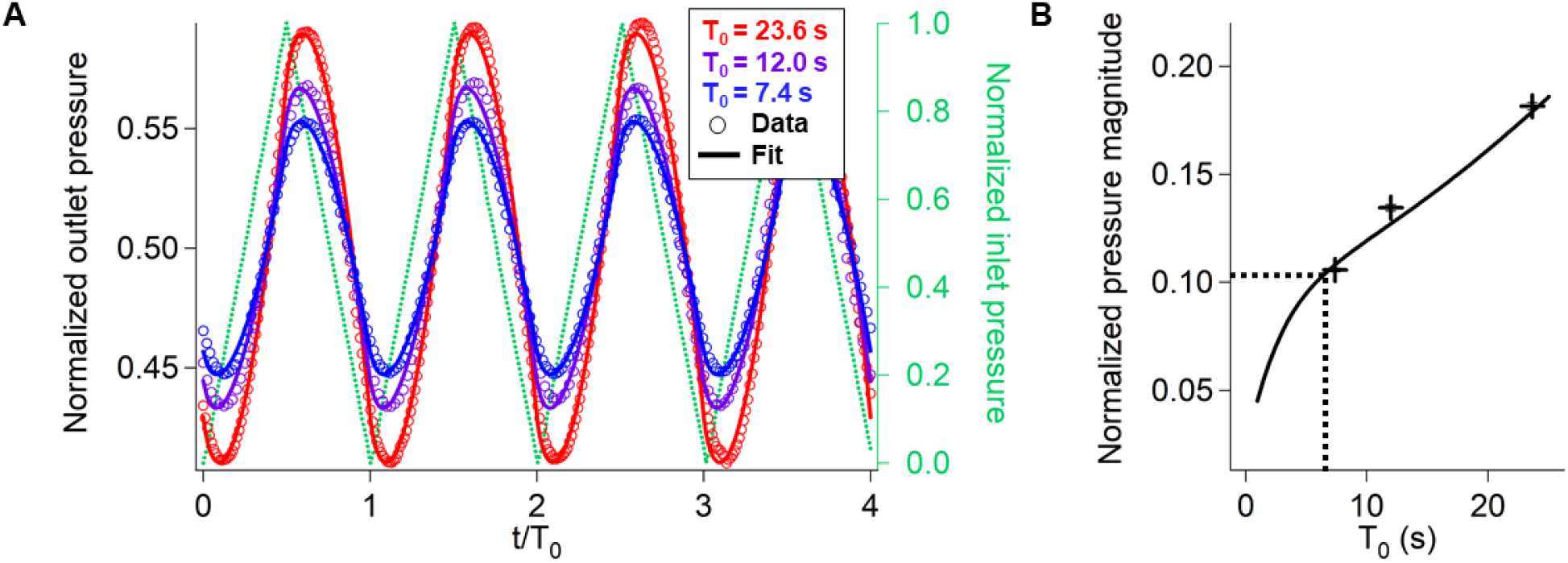
A) The pressure in the cavity is reported as a function of the time normalized by the period of the triangular wave input signal (green dashed line). The blue, purple, and red datapoints correspond to the signals collected for periods of 7.4, 12.0, and 23.6 s, respectively. The corresponding solid lines are fits with the linear poroelastic model and one set of parameters for the P-wave modulus and permeability of 1.7 kPa and 7 × 10^-14^ m^2^. B) The outlet pressure magnitude (crosses) increases with the period of the signal, in good agreement with the linear model shown by the solid line. The dashed lines show the period for an output pressure of 100 Pa (10% of 1000 Pa).

The analytical solution of the 1D linear poroelastic model defined in Eq. (4) exactly matched the outlet pressure variations for the three periods using one set of parameters for the P-wave modulus and permeability of (1667 ± 293) Pa and (6.5 ± 1.0) × 10^-14^ m^2^, respectively (solid lines in Fig. 2A). The consistency of the 1D model for the semi-circular geometry of this system (Fig. 1B) was already acknowledged in our previous report, in which we used step-pressure inputs instead of periodic triangles [21]. This consistency was essentially explained by the attachment of the gel at the bottom and lateral sides of the chip that force an essentially vertical pressure gradient. In line with this comment, the values of the P-wave modulus and permeability measured in this study were in good agreement with the average estimates of 1005 Pa and 6.8 × 10^-14^ m^2^, respectively, of our previous report.

Having established a quantitative match between the model and our data, we focused on optimizing the operation of this technology for monitoring the maturation of poroelastic materials. This optimization involved achieving a measurable pressure signal amplitude while minimizing the actuation period to enable characterizations on minute timescales. Given the sensor’s precision of 2.5 Pa (see methods), we considered that a signal forty times greater of 100 Pa was appropriate for quantitative recordings. For an input pressure of 1000 Pa, this amplitude is attained for a period of ∼6 s (dashed lines in Fig. 2B). Therefore, we conducted our study at a constant period of 6.7 s and averaging the pressure response over 10 cycles (*i*.*e*., for a total time of ∼1 minute).

### Fitting collagen gel responses with porohyperelastic models

We then examined the response of native collagen gels and identified distinct differences compared to fixed cross-linked collagen gels. We recorded the output pressure for 10 periods with 20 measurements per period, and combined the data into a master response curve for one single period (red dots, Fig. 3A). The output pressure was no longer centered at the average input pressure of 500 Pa but instead shifted to approximately 700 Pa. This shift indicates that the gel remained in compression (*P*(*H, t*) > *P*(0, *t*)) for a longer duration than in extension ((*P*(*H, t*) < *P*(0, *t*)). Such an output pressure offset was previously observed under sinusoidal wave pressure actuation [22] and attributed to the tension-compression asymmetry of the permeability of collagen gels. At steady state, the volume of fluid injected into the gel must equal the volume expelled from it over one period. Since permeability is higher in tension than in compression, this balance is achieved by spending more time in extension than in compression, which explains the pressure offset. In turn, this mechanism highlights the importance of accounting for the nonlinear properties of native collagen gels to accurately reproduce their responses.

**Figure 3:**
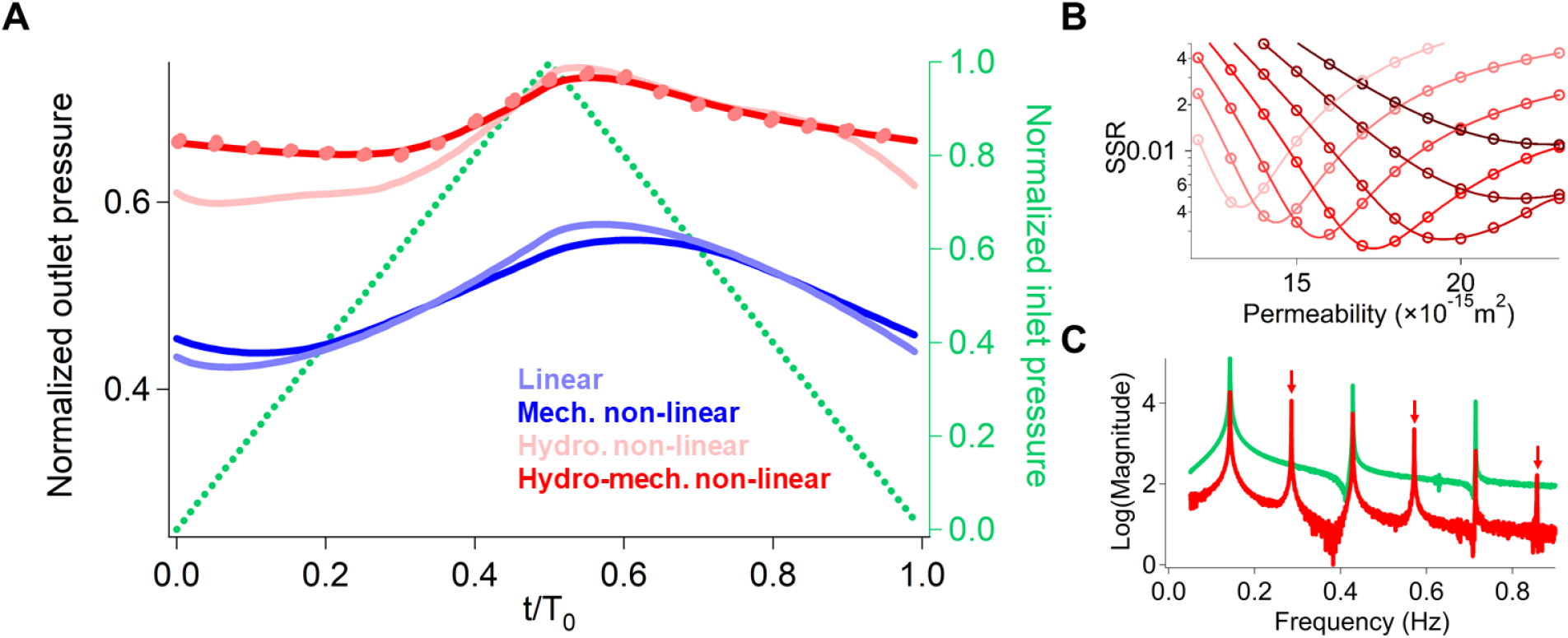
A) COMSOL simulations of the normalized pressure response as a function of the time normalized by the actuation period for M = 230 Pa and κ = 1.7×10^−14^ m^2^ (i) with linear poroelastic properties (blue curve), (ii) with non-linear mechanical behavior (dark blue curve), (iii) with non-linear permeability (light red curve), and (iv) with non-linear mechanics and non-linear permeability (dark red curve). The experimental response of native collagen gels is reported with red points. B) Sum of square residuals by comparison of the experimental data with the simulations for the p-wave moduli ranging from 200 Pa (light red) to 260 Pa (dark red) and permeabilities ranging from 1.2×10^-14^ m^2^ to 2.3×10^-14^ m^2^. C) Fourier Transform analysis of the inlet pressure (in green) and outlet pressure with native collagen (in red). Even frequencies are marked with red arrows.

We upgraded the poroelastic model with non-linear elasticity and/or non-linear permeability using the compressible neo Hookean and Kozeny-Carman models, respectively (see Eq. (7) and (8)). We could not solve these nonlinear models analytically, and focused on the implementation of finite element simulations (Fig. 3A). We first incorporated non-linear mechanical properties and compared it to the linear response described in the previous section (light blue curve in Fig. 3A). The strain-stiffening behavior tended to smooth out the pressure variations in the cavity but did not cause any offset in the response (dark blue curve in Fig. 3A). We then implemented a model with linear elasticity and a quadratic variation of the permeability as a function of the deformation (Eq. (7)). This model enabled us to fit the offset (light red curve in Fig. 3A), but it did not provide a satisfactory fit to the data because the amplitude of the predicted signal was larger than the experimental readout. We achieved a good fit by combining strain-stiffening with the quadratic variation of the permeability because this model allowed us to reduce the amplitude of the output pressure variations (dark red curve in Fig. 3A, respectively). The fitting of the data allowed us to infer the P-wave modulus and the permeability of 230 ± 10 Pa and 1.7 ± 0.1 ×10^−14^ m^2^, respectively. Note that the minimization of the residue between the data and the model yielded a single extremum in the parameter space (Fig. 3B). These values fell within the same range as those measured for cross-linked collagen gels but were lower. The seven-fold decrease in elasticity was anticipated, as cross-linking agents are known to stiffen collagen gels [25]. However, the change in permeability was more challenging to interpret. Linear poroelasticity analysis revealed a consistent five-fold decrease in compression between cross-linked and native collagen gels, yet the values remained comparable in tension. It is important to note that porohyperelasticity offers an integrated analysis of collagen gel responses across various actuation ranges, where linear models produced stress-dependent parameters.

Further, we performed Fourier Transform analysis of the response of collagen gels (Fig. 3C), which showed the presence of even and odd frequencies (red curve in Fig. 3C) whereas even frequencies were only excited by the periodic triangle actuation (gray curve in Fig. 3C). This result was consistent with the porohyperelastic model, in which the quadratic dependence of the non-linear permeability generates modes at double the frequency of the input signal in Fourier space. Interestingly, this spectrum was obtained disregarding the offset of the outlet pressure, which nevertheless provided the first hint to the non-linear permeability of collagen gels. Furthermore, the simulation only built on non-linear mechanics did not produce odd frequency harmonics in the Fourier spectrum (not shown), implying that the response of the native collagen slab was primarily dictated by its non-linear permeability, as predicted by the Kozeny-Karman model.

### Tissue remodeling by senescent fibroblasts

Using our methodology for the parametrization of a porohyperelastic model, we then investigated the effect of senescent fibroblast cells on the properties of collagen gels. We applied the same periodic triangular pressure actuation with a period of 6.7 s, and recorded the pressure response in the air cavity for 10 to 20 periods (i.e., approximately 2 minutes). We performed measurements at day 0 for the reference response of native collagen gels, and after 4, 7, and 10 days of maturation in the presence of cells. While we did not observe any change of the response for an unseeded gel (data not shown), the pressure was consistently shifted upon cell incubation (datapoints in Fig. 4A). We observed a gradual onset of the average pressure in the cavity from 700 to 850 Pa, along with a reduction in the pressure oscillation amplitude by half, from approximately 100 Pa to 50 Pa. The same porohyperelastic model allowed us to reproduce the four curves each with two fitting parameters *κ*_0_ and *M*_0_, demonstrating the relevance and strength of our methodology to investigate the maturation of biological tissues.

**Figure 4:**
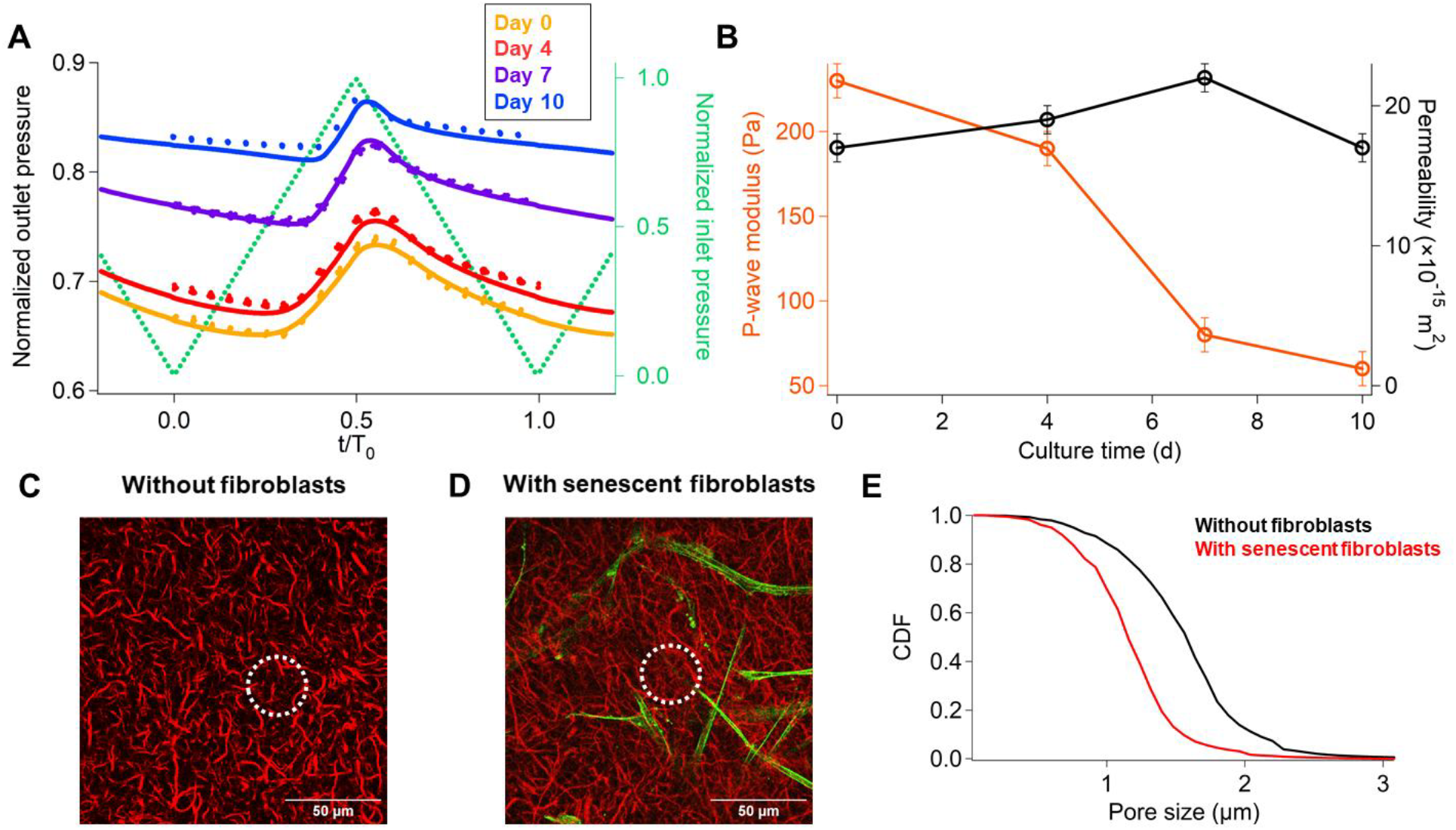
A) Pressure responses (circles) and fits (solid lines) for collagen gel seeded with senescent fibroblasts at different culture times. The actuation triangular pressure inlet is shown in green. B) Fitted P-wave modulus and permeability over culture time. C) Immunostaining of native collagen (in red) without and D) with senescent fibroblasts (in green) shows an unaltered *vs*. a distinct reorganization of the collagen fibers secondary structure (white dashed circles), respectively, indicative of senescence-induced remodeling after 4 days of culture. E) Cumulative density function (CDF) of the pore radius in collagen without (black) and with (red) senescent fibroblasts after image analysis. (N = 3 for each condition, and 40 images of each 70 µm-stack have been analyzed).

Data fitting shows that the permeability of collagen gels remained constant at 18.8 ± 2.4 × 10^-15^ m^2^ and equal to that of the native material (Fig. 4B). Conversely, the tissue appeared to become softer, as the P-wave modulus dropped by a factor of 3.8 after 10 days (this effect was consistently observed across independent batches of tissue fabrication). It may appear surprising that the model predicts a constant permeability despite the onset of the pressure offset during the maturation. In fact, the enhanced deformability of the tissue enhances the variation of the permeability in tension *vs*. compression, forcing a shift of the pressure offset in the cavity to balance influx and outflux in the material. We finally sought to gather evidence about the ECM reorganization induced by senescent fibroblasts. We performed immunostaining with an antibody targeting collagen fibers in the gel seeded or not with fibroblasts (Fig. 4C-D, respectively). Confocal micrographs showed that the total intensity signal of the collagen fibers was roughly constant in collagen gels or senescent tissues, in agreement with the fact that collagen production is severely reduced in senescent fibroblasts [31]. Moreover, the data suggested that senescent fibroblasts remodeled the collagen gel microstructure, as indicated by an increased fiber density and more branching. This remodeling occurred throughout the entire tissue, not just near the cells, and could be quantitatively assessed using image-based porosimetry. The apparent pore size decreased from 1.6 to 1.2 µm in the senescent tissue (Fig. 4E). However, the relationship between the microstructure and the poromechanical properties of the network remains unclear. Assuming the total collagen concentration stays constant, increased branching implies thinner fibers, which could reduce their elastic modulus and soften the tissue [32]. Yet, the tissue’s constant permeability suggests that the pore size remains unchanged, contradicting our structural data. It is important to note that permeability is associated with a length scale 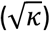 of approximately 140 nm, which is below the resolution limit of confocal microscopy. This raises questions about the relationship between porosity and poromechanical measurements, beyond the evident network reorganization.

## Discussion

This study introduces a microfluidic platform that enables the real-time assessment of poromechanical properties in engineered tissue models, offering insights into the impact of stromal cells on virtually any hydrogel. By combining periodic pressure actuation with advanced modeling, we provide a robust method to probe the dynamic interplay between fluid transport and tissue deformation under nonlinear mechanical and hydrodynamical conditions. Our findings demonstrate that senescent fibroblasts induce a significant reduction in the P-wave modulus of collagen gels over time, indicative of tissue softening, while hydrodynamic permeability remains constant. This approach not only advances the field of tissue mechanics but also sheds light on the complex mechanisms of tissue remodeling during aging.

A key advantage of our platform is the integration of periodic actuation to interrogate both elastic and porous contributions to tissue mechanics. This approach allows us to capture the non-linear mechanical behaviors of collagen gels, including strain-stiffening and deformation-induced permeability changes. By coupling experimental measurements with a detailed computational framework based on compressible Neo-Hookean mechanics and the Kozeny-Carman permeability relation, we provide a physiologically relevant model of the ECM. Moreover, the on-chip integration of the tissue model minimizes contamination risks and enables long-term monitoring, making this system ideally suited for studying age-related tissue changes *in vitro*.

The observed reduction in the P-wave modulus can be attributed to several potential mechanisms related to the activity of senescent fibroblasts. Immunostaining of the collagen gels revealed significant remodeling of the ECM structure, that may be induced by the secretion of matrix metalloproteinases by senescent fibroblasts, which degrade collagen and other ECM components. This enzymatic activity could contribute to the progressive softening of the collagen matrix.

The implications of these findings extend beyond tissue mechanics. A softer ECM may influence the behavior of mechanosensitive cells, such as immune cells, fibroblasts, and epithelial cells, potentially exacerbating inflammatory and degenerative processes in aged tissues. Looking ahead, investigating whether the softening of the ECM can be reversed through the removal of senescent fibroblasts or inhibition of their secretory phenotype could provide insights into potential therapeutic strategies. Furthermore, the platform could be used to test ECM-stabilizing drugs or senolytic agents to evaluate their efficacy in restoring tissue mechanics and countering age-related degeneration.

In conclusion, this study highlights the profound impact of senescent fibroblasts on ECM poromechanics, revealing a significant reduction in tissue elasticity despite stable permeability. The developed platform represents a powerful tool for dissecting the complex interplay between cellular activity and ECM remodeling, offering valuable insights into the mechanisms underlying tissue aging and degeneration.

## Materials and method

### Cell culture

All cells were maintained at 37 °C in a humidified atmosphere of 5% CO2/95% air. Primary human skin fibroblasts HCA2 (MJ90) were previously isolated from neonatal foreskin by the laboratory of Judith Campisi (University of California San Francisco, USA). Fibroblasts were cultured in high glucose Dulbecco’s Modified Eagle’s Medium (DMEM) supplemented with 10% fetal bovine serum (FBS; Biosera, Nuaille, France) and 1% penicillin–streptomycin (pen–strep; FUJIFILM Wako Pure Chemical Corporation, Osaka, Japan). For passaging, cells were rinsed once with 1× Dulbecco’s phosphate buffered saline (−) (PBS; FUJIFILM Wako Pure Chemical Corporation), incubated with 0.25 w/v% trypsin-5.3 mmol L−1 EDTA-4Na solution (trypsin; FUJIFILM Wako Pure Chemical Corporation) for up to 5 minutes at 37 °C, and harvested in culture medium. To induce senescence of skin fibroblasts, 70– 80% confluent early passage HCA2 were irradiated in culture medium with 10 Gy of γ-ray by using a MBR-1520R-3 (Hitachi, Tokyo, Japan). Senescent cells were then selected by culturing for 14 days with DMEM containing 10% FBS and 100 nM BI2536 (Plk1 inhibitor; Cayman Chemical, MI, USA), and senescence phenotype was first checked as previously described in ref. [17].

### Chip preparation

The polydimethylsiloxane (PDMS)-based chips (25 mm × 25 mm × 5 mm: width × length × height) and their fabrication method were previously described. [33] To prepare a 3D tissue with senescent skin fibroblasts, a PDMS chip was treated with air plasma (basic plasma cleaner; Harrick Plasma, Ithaca, NY, USA) for one minute and sterilized by UV-light under a cell culture hood. A ∅ 200 μm acupuncture needle (No. 02, 0.20 mm × 30 mm, J type; Seirin, Shizuoka, Japan) was coated with 1% BSA in PBS, dried, and also sterilized by UV-light. The chips were treated with 50 μl of 2.5% glutaraldehyde for 1 min, then thoroughly rinsed with water, and dried. The collagen solution was subsequently prepared on ice by mixing Cellmatrix Type I-P (pepsin solubilized) solution (Nitta Gelatin, Japan), 10× Hanks’ buffer, and 10× collagen buffer (volume ratio, 8:1:1) following the manufacturer’s protocol. Senescent MJ90 fibroblasts were harvested and resuspended at a density of 200 000 cells per mL in the collagen solution. Fibroblasts-containing collagen solution was introduced into the PDMS chip chamber and the BSA-coated acupuncture needle was inserted through the PDMS channel. The chip was incubated at 37 °C for 30 min to enable gelation of collagen solution, and the acupuncture needle was withdrawn to leave an empty channel in the collagen gel. Warm medium (DMEM supplemented with 10% fetal bovine serum and 1% penicillin–streptomycin) was then added and the 3D model was cultured at 37 °C with medium change every 2 days.

### Elasticity and permeability measurements

The collagen gels embedded within a silicone chip was connected to a pressure controller (Fluigent MFCS) and a pressure sensor (Merit Sensor LP series) by means of 3D printed mechanical pieces (Elegoo Mars 3, water washable resin). Acquisition was done through a dedicated LabVIEW interface allowing the control of the inlet pressure while recording the outlet signal with a digital multimeter (Agilent, 34401A). The resulting data were fitted by either coding the analytical solution in Spyder and using the “curve_fit” function to retrieve *M* and κ, or comparing the simulation responses with the experimental data, minimizing the sum of square roots.

### COMSOL simulations

Simulations were run with COMSOL Multiphysics 6.2 using the poroelastic module, which allowed coupling of the Darcy’s law and the solid mechanics modules. The 1D geometry consisted in a line of 0.8 mm height. An inlet pressure *P*_*M*_ was imposed at *x* = 0, creating a double boundary condition defined as a pressure in the Darcy’s law module and a boundary load in the solid mechanics module. We used the “boundary ODEs (ordinary differential equations) and DAEs (differential-algebraic equations)” module to solve the boundary condition in the air cavity, and compute the integral of the flow for each time point. Lastly, a weak contribution and an auxiliary dependent variable were used in the solid mechanics physic to map the computed Jacobian value of 1D deformation to a new variable. Finally, the Young Modulus, Poisson ratio and permeability were defined as a function of this new variable in order to map the stress and hydrodynamic pressure.

### Immunofluorescence staining

Immunofluorescence confocal microscopy was performed with the LSM 700 confocal microscope (Carl Zeiss) equipped with a 40× water immersion objective (numerical aperture of 1.2) and using a pinhole of 1 Airy unit. The samples were fixed with 4% (w/v) paraformaldehyde in PBS (FUJIFILM Wako Pure Chemical Corporation) for 30 minutes, and permeabilized with 0.5% Triton X-100 in PBS for 10 minutes at 25 °C. Blocking was performed by incubating overnight at 4 °C with blocking solution (1% (w/v) BSA in PBS). Samples were then incubated with Alexa Fluor 488 phalloidin (Thermo Fisher Scientific) for actin staining. The microstructure of native or cross-linked collagen gels was observed after incubation with the conjugate of mouse monoclonal anti-collagen I antibody (ab6308, Abcam, Cambridge, UK) and Alexa Fluor 488-labeled donkey antibody agonist mouse IgG diluted at 1:100 in blocking solution for 4 hours, and then rinsing the gel with phosphate buffer.

### Confocal images analysis

Z-stack images of collagen fibers were first processed using ImageJ. 3D Gaussian blur filter (Radii x=2, y=2, z=1 in pixels) was applied before applying an individual threshold filter to each image slice to isolate the main collagen fibers into a 3D binary file. These binary files were further analyzed using a python model named Porespy, [34] a collection of image analysis tools to extract information from 3D images of porous structures. We used the porosimetry function to extract the pore size cumulative distribution function of the collagen binary files.

## Acknowledgment

The authors acknowledge Eri Otsuka and Takanori Sano (The University of Tokyo) for their technical assistance.

## Author contribution

Conceptualization: JC, AB

Methodology: JC, AB, JOM

Investigation: JC, TQ

Software: LJ

Materials: MN

Visualization: JC, DA, SN

Supervision: JC, PC, AB, YTM

Writing—original draft: JC

Writing—review & editing: JC, AB

## Data and materials availability

All data are available in the main text or the supplementary materials.

## SUPPLEMENTAY MATERIAL

**Figure S1:**
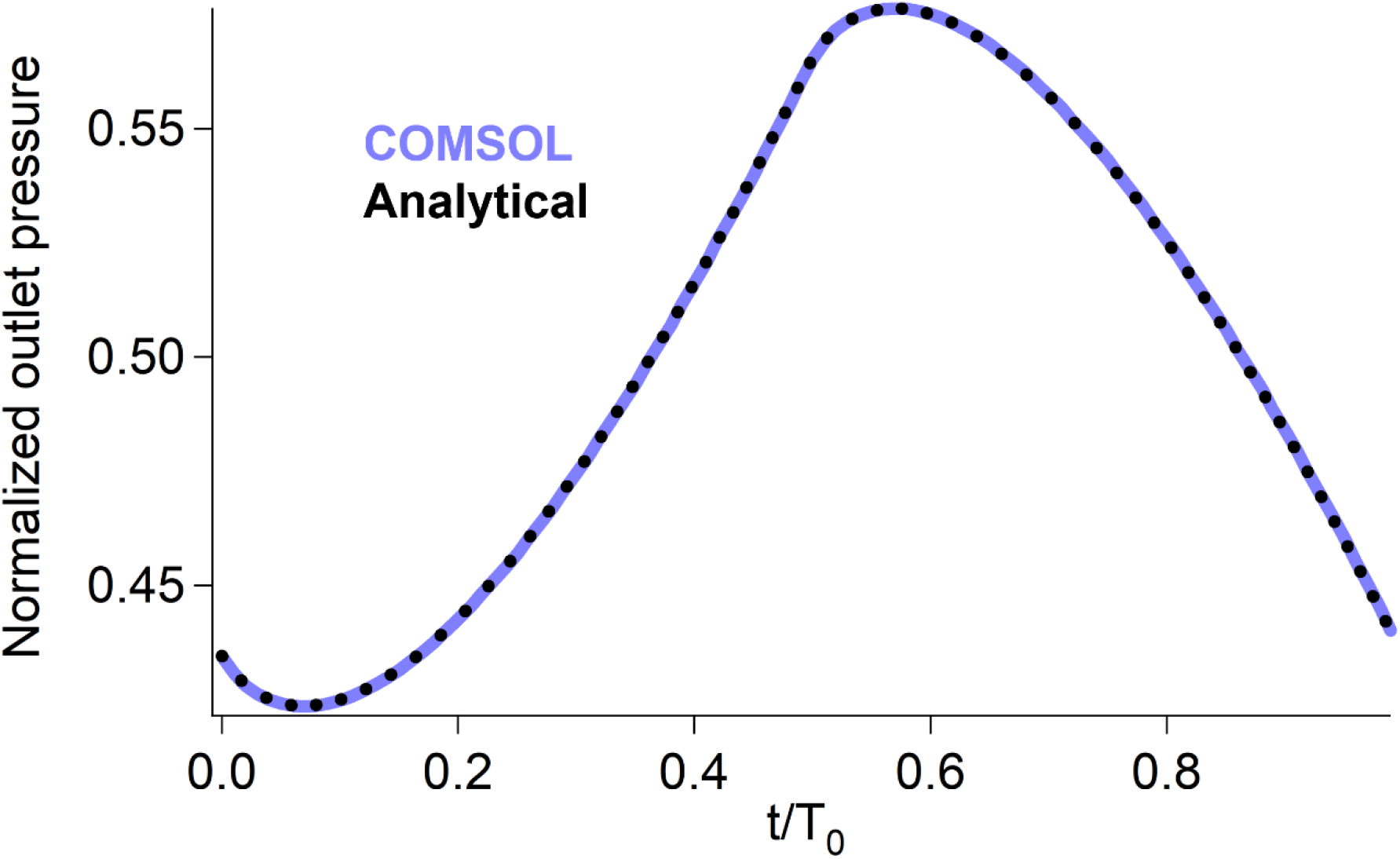
comparison of the 1D COMSOL solution with the analytical solution. We consider here a material with a permeability of 1.7×10^-14^ m^2^ and a p-wave modulus of 230 Pa for a triangle inlet pressure period of 6.7 s.

### 1D model

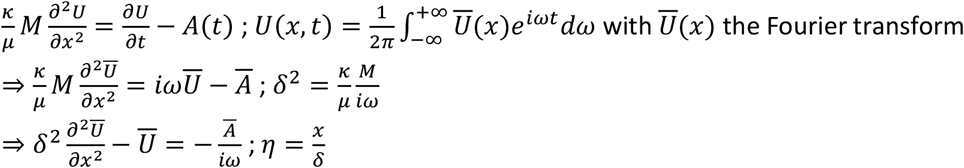

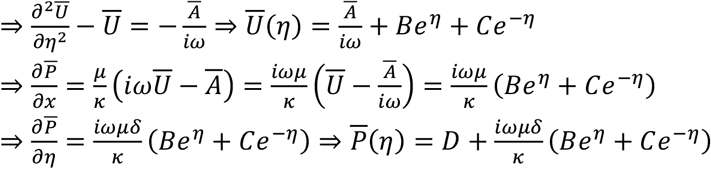

#### Boundary conditions

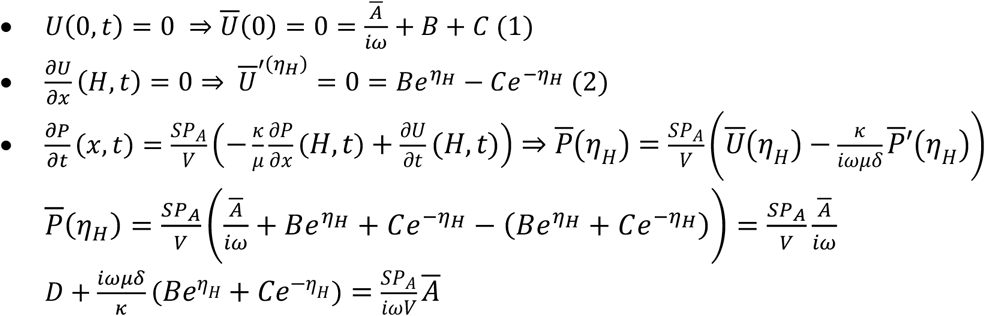

From (2), we have 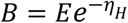 and 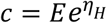

From (1), we have 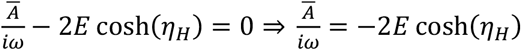

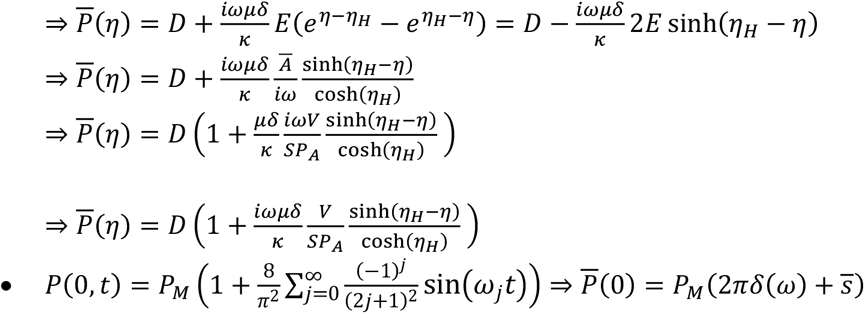

with 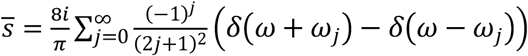 and 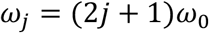

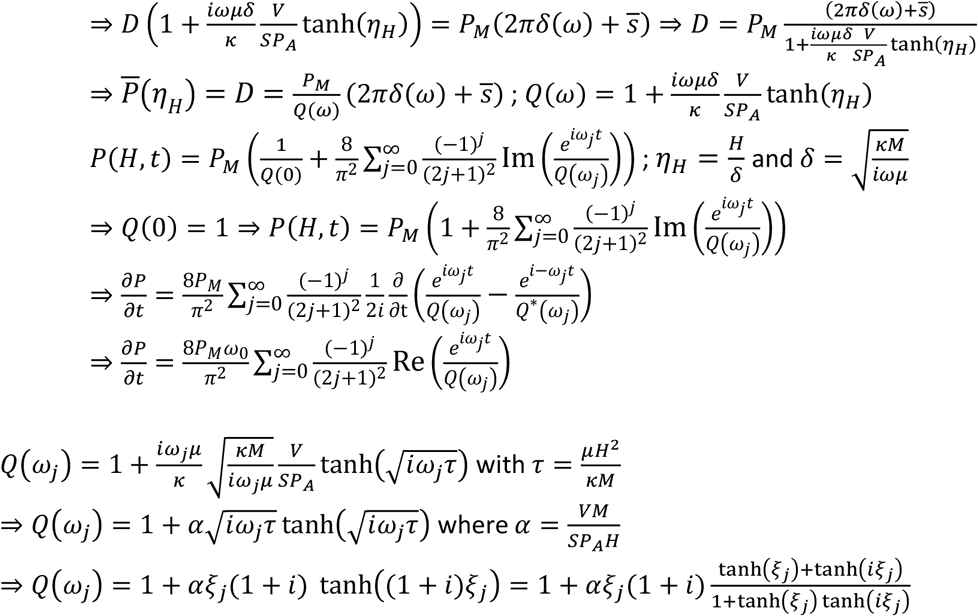

with 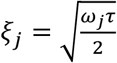 and tanh(i*ξ*_*j*_) = *i*tan(*ξ*_*j*_)

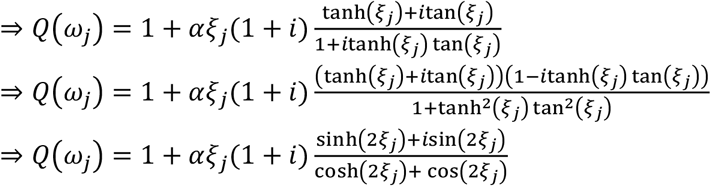

We now define 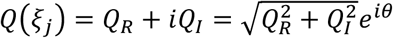 where 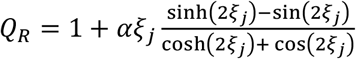 and 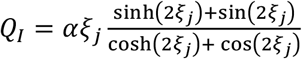

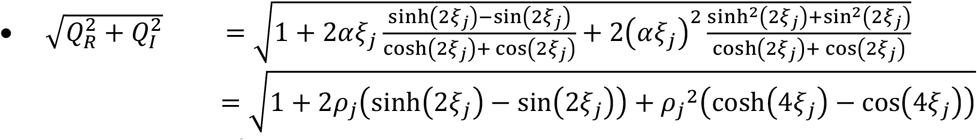

where 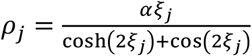

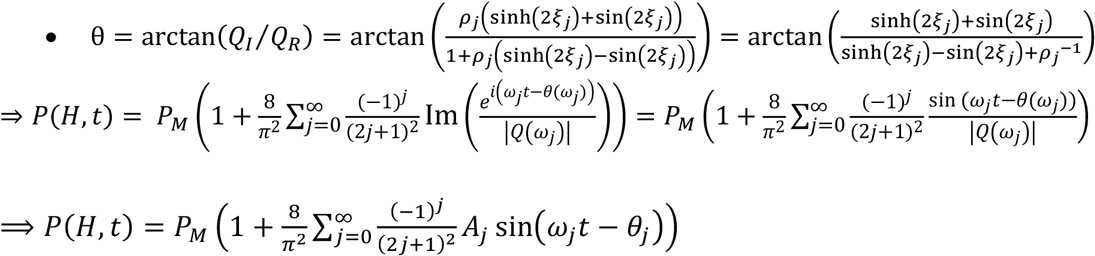

with 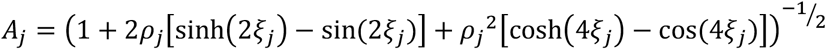 and

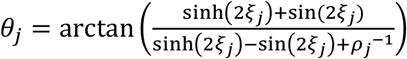

